# Plasticity in gene expression facilitates invasion of the desert environment in house mice

**DOI:** 10.1101/2020.02.10.939231

**Authors:** Noëlle K. J. Bittner, Katya L. Mack, Michael W. Nachman

## Abstract

Understanding how organisms adapt to new environments is a key problem in evolution, yet it remains unclear whether phenotypic plasticity generally facilitates or hinders this process. Here we studied the evolved and plastic responses to water stress in lab-born descendants of wild house mice (*Mus musculus domesticus*) collected from desert and non-desert environments. Using a full sib design, we measured organismal phenotypes and gene expression under normal (hydrated) and water stressed (dehydrated) conditions. After many generations in the lab, mice from the desert consumed significantly less water than mice from other localities, indicating that this difference has a genetic basis. Under water stress, desert mice lost less weight than non-desert mice, and desert mice exhibited differences in blood chemistry related to osmoregulatory function. Gene expression in the kidney revealed evolved differences between mice from different environments as well as plastic responses between hydrated and dehydrated mice. Desert mice showed reduced gene expression plasticity under water stress compared to non-desert mice. Importantly, the non-desert mice generally showed shifts towards desert-like expression under water stress, consistent with adaptive plasticity. Finally, patterns of gene expression identified several candidate genes for adaptation to the desert, including *Aqp1* and *Apoe*. These findings provide evidence for local adaptation in a recently introduced species and suggest that adaptive plasticity may have facilitated the colonization of the desert environment.

## Introduction

Understanding the origin and genetic architecture of complex traits associated with local adaptation is a central goal of evolutionary biology. One ongoing debate concerns the extent to which phenotypic plasticity may facilitate or constrain adaptation to new environments (Baldwin, 1896; Price, Qvarnström, & Irwin, 2003; Ghalambor, McKay, Carroll, & Reznick, 2007; Levis & Pfennig, 2016). For example, adaptive plasticity, defined as an environmentally induced phenotypic change that brings individuals closer to the local optimum, may enable organisms to invade new environments. Subsequent genetically encoded changes in the same direction as the plastic changes may then accrue, bringing individuals even closer to the optimum, as seen for coloration in lizards living on dark substrates (Corl et al., 2018). Conversely, plastic changes may be non-adaptive, moving individuals farther from the local optimum. In such cases, selection is expected to favor genetic changes underlying phenotypes that go in the opposite direction of the plastic change and thereby bring the individual closer to the optimum. This pattern of non-adaptive plasticity is seen for gene expression changes in guppies reared in the absence of predators (Ghalambor et al., 2015). Which of these two outcomes is most likely remains unclear and may depend both on the phenotype in question and the environmental heterogeneity to which populations have been exposed (e.g. Huang & Agrawal, 2016).

House mice (*Mus musculus domesticus*) provide an opportunity to study plastic and evolved changes in the context of adaptation to novel environments. House mice are native to western Europe but were recently introduced to the Americas with European colonization, approximately 400-600 generations ago (Phifer-Rixey & Nachman, 2015). In this short time, they have colonized a wide variety of different environments. In eastern North America, house mice show strong evidence of local adaptation for several complex phenotypes such as body size, activity, and nest-building behavior (e.g. Lynch, 1992; Mack, Ballinger, Phifer-Rixey, & Nachman, 2018; Phifer-Rixey et al., 2018).

In the Sonoran Desert of North America, house mice must contend with low to seasonally-absent water as well as extreme heat. Although house mice are human commensals, they frequently live in sheds, grain storage areas, barns, and other habitats where they are not well shielded from the environment. They can also live in situations where they are not associated with humans (Sage, 1981). House mouse urine concentration, a metric often associated with specialization to xeric environments, is very high (Haines et al., 1973) and falls within the range of many known desert specialists (Beuchat, 1990). Previous experiments of wild mice brought into the lab have found that mice can survive beyond 14 days without access to free water on a diet of dried seeds (Koford, 1968). In other experiments with varying relative humidity, mice survived up to 41 days without free water (Haines & Schmidt-Nielsen, 1967).

Previous studies have examined the behavioral, morphological, and physiological adaptations that allow desert mammals to persist under xeric conditions (reviewed in Schmidt-Nielsen, 1964; Donald & Pannabecker, 2015). Recent work has also begun to identify some of the genes associated with these phenotypes (Giorello et al., 2018; MacManes, 2017; Marra et al., 2014; Wu et al., 2014). Far less is known about the role of phenotypic plasticity in the context of desert adaptation. Since all mice, including those living in more mesic environments, occasionally go through periods of water stress, selection may have favored plastic responses that enable mice to survive periods of water shortage (i.e. adaptive plasticity). Here we are interested in whether the evolved differences between mice from more mesic environments and mice from more xeric environments mirror the plastic responses seen within populations, both for organismal level phenotypes and for gene expression in the kidney. Changes in gene expression in the kidney may also help identify candidate genes for adaptation to xeric conditions.

To assess the contribution of plastic and evolved changes to a xeric environment, we studied lab-born descendants of wild mice from two populations, one from Edmonton, Canada and one from Tucson, Arizona. While neither population experiences high precipitation, the annual precipitation in Edmonton is 57% more than in Tucson, and these two locations differ dramatically in average temperature. We address four primary questions. (1) Do progeny of wild mice from these two populations exhibit phenotypic differences when reared in a common laboratory setting, indicating that the phenotypes have a genetic basis? (2) Do these same phenotypes exhibit plastic (non-genetic) changes when mice are exposed to water limitation? (3) Are plastic changes generally in the same or opposite direction as the evolved changes? (4) Do gene co-expression networks identify sets of genes and corresponding phenotypes that underlie adaptation to xeric conditions? We find that in a common environment, Tucson mice drink less water than Edmonton mice and differ in blood chemistry and gene expression in the kidney. These same traits exhibit significant plasticity when mice are fully hydrated compared to mice under water stress, and evolved differences are generally in the same direction as plastic differences, both for gene expression and for organism-level phenotypes. Finally, co-expression networks identify groups of genes that likely underlie adaptation to xeric conditions. These findings suggest an important role for adaptive plasticity in the colonization of the desert environment.

## Materials and Methods

### Mice

To assess whether mice from the Sonoran Desert differ in water consumption compared to mice from other habitats, we used wild-derived mouse lines developed in our lab from a range of localities in the Americas. Wild house mice were caught from five populations in different habitats and used to create inbred lines through sib-sib mating over multiple generations. The five localities were Tucson, AZ, USA (TUC); Edmonton, Alberta, Canada (EDM); Gainesville, FL, USA (GAI); Saratoga Springs, NY, USA (SAR); and Manaus, Amazonas, Brazil (MAN). In nearly all cases, lines were established from unrelated individuals. Lines were maintained in the laboratory for 6-19 generations on a diet of standard mouse chow. All mice were handled in accordance with a UC Berkeley Animal Care and Use protocol (protocol AUP-2016-03-8548-1).

### Measuring water consumption

To quantify differences among populations, we measured water consumption over 72 hours in 163 adult males representing 45 different inbred lines (Table S1). In total, 40 individuals were measured from Tucson, 24 from Gainesville, 48 from Edmonton, 23 from Saratoga Springs, and 28 from Manaus. Mice were between 90 and 200 days of age and were housed singly in cages at 23°C with a 10 hour dark and 14 hour light cycle on standard Teklad Global rodent chow (18% protein, 6% fat). Body weight and amount of water consumed after 72 hours were recorded, and relative water consumption (RWC) was calculated (grams of water consumed per gram of mouse). Relative water consumption was used as a metric due to the population level variation in body weight.

### Measuring phenotypic plasticity

To study evolved and plastic responses to xeric conditions, we chose one wild-derived inbred line each from Tucson and Edmonton [Tucson: TUSA4xA8 (TUCC/Nach), Edmonton: TAS111x165 (EDMA/Nach)]. These lines were chosen because they showed large differences in water consumption as well as little variance within lines. Male littermates from these lines were assigned at random to either control or water restriction treatments and housed individually post-weaning with water *ad libitum*. After 90 days of age, mice were weighed and phenotyped for relative water consumption. Following phenotyping (average age = 99 days), mice assigned to the water restriction treatment (Edmonton n=11, Tucson n=7) were restricted from all water consumption for 72 hours. Mice assigned to the control treatment (Edmonton n=10, Tucson n=6) were maintained with water *ad libitum*. All mice were weighed every 24 hours and monitored for markers of drastically declining health. All animal care was conducted in accordance with procedures approved by the UC Berkeley Animal Care and Use Committee (protocol AUP-2017-05-9940). After sacrifice with isofluorane, left kidney, liver, and caecum were immediately removed and stored in RNAlater according to manufacturer’s instructions. The right kidney was weighed and stored in 10% neutral buffered formalin for morphological analysis at the UC Davis Comparative Pathology Laboratory. Five hydrated mice per population (ten mice total) were phenotyped for renal cortex thickness and papilla thickness. Blood was extracted from the heart and body cavity using a syringe and centrifuged in BD SST Microtainer tubes to extract serum. Levels of blood urea nitrogen (BUN), total protein, creatinine, chloride, potassium, and sodium levels were analyzed at the UC Davis Comparative Pathology Laboratory for 20 mice (five individuals per population per treatment). These serum solutes were measured to quantify kidney health and glomerular function in treated and control samples.

### mRNA library preparation and sequencing

RNA was extracted from half of a kidney preserved in RNAlater from twenty mice total, (five mice per population per treatment), using the MoBio Laboratories Powerlyzer Ultraclean Tissue & Cells RNA Isolation Kit. RNA libraries were prepared using the KAPA Hyper Prep Kit and then pooled and sequenced across two lanes of 100bp PE Illumina HiSeq4000 at the Vincent J. Coates Genomics Sequencing Center at UC Berkeley.

### mRNA read mapping and quantification of gene expression

Reads were trimmed for quality and adaptor contamination with Trimmomatic v0.36 (Bolger et al., 2014) and mapped to the *Mus musculus* reference genome (GRCm38/mm10) using STAR v2.6.0c (Dobin et al., 2013). Reads overlapping exons were counted using the program HTSeq 0.6.1 (Anders et al., 2015) to estimate per-gene mRNA abundance. We removed genes with a mean fewer than ten reads across samples from additional analyses. The R package DESeq2 (Love et al., 2014) was used to test for differential expression between (1) treatments within each population, and (2) populations within each treatment. Genes were retained as significant at a false-discovery rate of 5%.

### Gene co-expression analyses

We used standard protocols (Langfelder & Horvath, 2008) to perform a weighted gene co-expression network analysis on expression residuals for the 20 individuals to identify expression modules. We tested for associations between eigengenes (the first principle component of a module) and each of the nine phenotypes described (RWC, kidney weight, proportion of weight maintained, serum BUN, serum creatinine, serum total protein, serum chloride, serum potassium, and serum sodium) as well as population of origin and treatment group. We were able to assign genes membership to expression modules as well as position relative to the center of the module. Genes that are more central (i.e. those that have more connections with other genes in a module) are good targets for putative candidate genes related to phenotypes of interest, population of origin, or treatment group. Additionally, to identify genes that show differential co-expression between Tucson and Edmonton we used the program DGCA (Differential Gene Correlation Analysis) (McKenzie et al., 2016), which calculates the average change in correlation between the two lines across all gene pairs.

### Enrichment analyses

GO category enrichment on gene sets of interests were performed with GOrilla (Eden et al., 2009) by testing the foreground set against the background set of all genes expressed in the kidney. Phenotype enrichment tests were performed with modPhea (Weng & Liao, 2017) by comparing the foreground set against the background set of all genes expressed in the kidney.

## Results

### Relative water consumption is lowest in mice from the Sonoran desert

To determine whether mice from Tucson, Arizona exhibit lower water consumption compared to mice from other populations, we took advantage of a set of wild-derived inbred lines of mice from five localities across the Americas (Figure 1a). We assayed relative water consumption for 163 male mice representing 45 wild derived inbred lines over a 72-hour period from five founder populations (Tucson (TUC), Edmonton (EDM), Saratoga Springs (SAR), Gainesville (GAI), and Manaus (MAN). Mice from Tucson drank significantly less water than mice from any other population except Saratoga Springs (Median RWC: TUC: 0.34, SAR: 0.40, MAN 0.42, GAI: 43, EDM: 0.52)(Figure 1b). The greatest difference was seen between mice from Tucson and mice from Edmonton (Mann-Whitney *U*, *p* < 0.00001)(Figure 1b). For this reason, we chose to focus on comparisons between lines from these two populations in all subsequent analyses.

**Figure 1.**
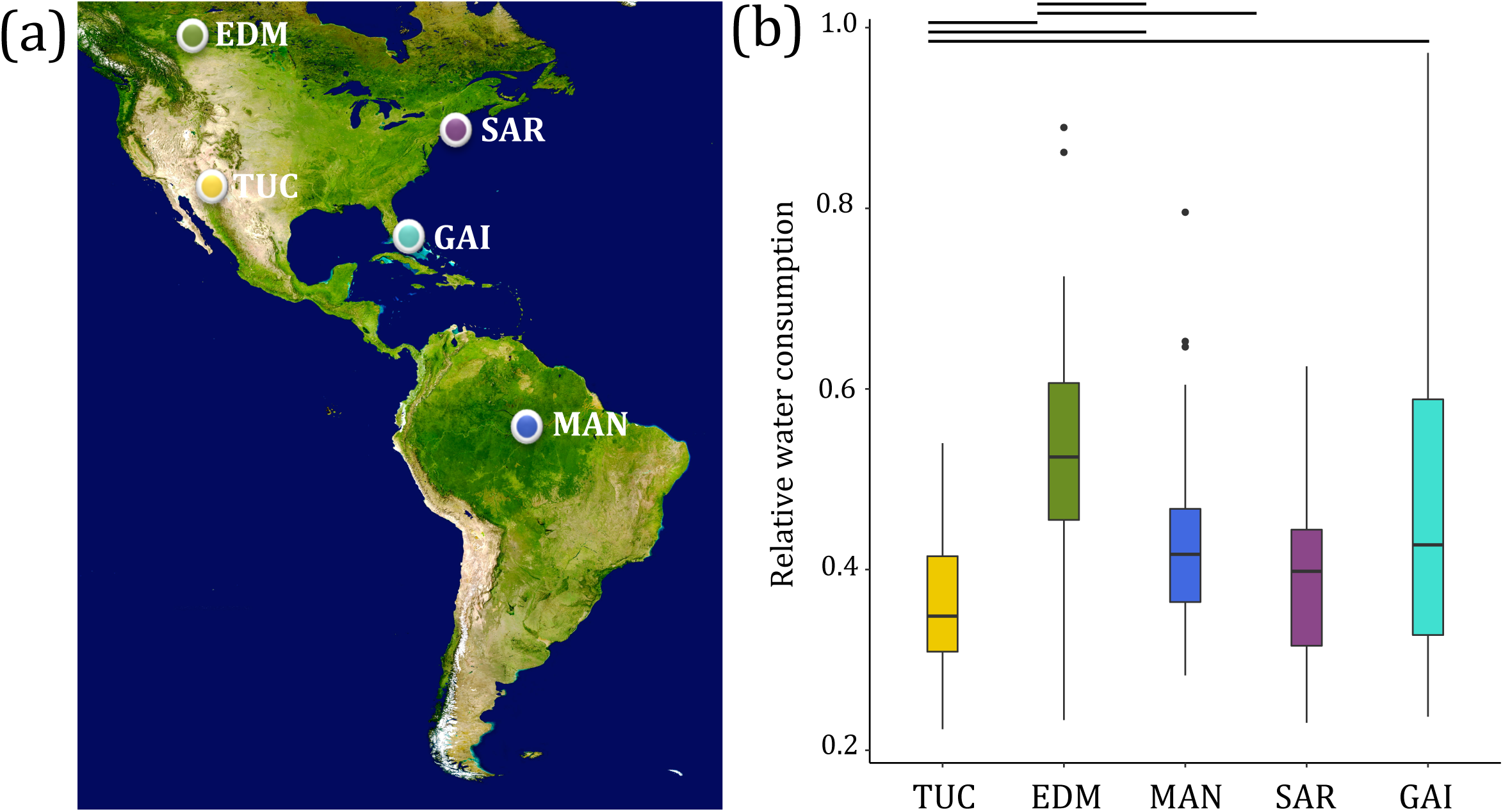
Relative water consumption in lab-born descendants of wild mice from different environments. (a) Sampling localities of wild-caught mice used to establish inbred lines in this study (map obtained from Google satelites.pro): Edmonton, Canada (EDM), Tucson, AZ (TUC), Gainesville, FL (GAI), Saratoga Springs, NY (SAR), and Manaus, Brazil (MAN). (b) Relative water consumption (g water consumed/ g mouse) in descendants of mice from different localities. Lines indicate comparisons that are significant (p < 0.05; Mann-Whitney U tests). Vertical lines denote 1.5 * the interquartile range.

### Evolved and plastic phenotypic differences associated with xeric conditions

Water consumption is a complex trait with many factors contributing to the ultimate phenotype. To further characterize this phenotypic variation, we compared the inbred line from Tucson with the lowest average water consumption (TUCC/Nach) to the inbred line from Edmonton with the greatest average water consumption (EDMA/Nach).

First, we compared mice from these two lines under standard (hereafter, “hydrated”) conditions. In addition to the difference in water consumption seen between mice from these lines (Figure S1), we found that hydrated mice from Tucson and Edmonton showed several phenotypic differences related to fluid consumption and homeostasis in a standard laboratory environment. Edmonton mice had heavier kidneys relative to their body weight than Tucson mice (*p* = 0.00015)(Figure S2), but did not show significant differences in two aspects of gross morphology: renal cortex thickness nor the ratio of papilla to cortex thickness (Figure S3). The relative thickness of the papillae in the medulla is correlated with urine concentrating ability; thicker medullas are often associated with animals inhabiting more arid climates (Al-kahtani et al., 2004). Anecdotally, mice from Tucson appeared to produce far less urine than mice from Edmonton, consistent with many desert rodents, but this was not measured in this study due to difficulty in obtaining urine from Tucson mice. In blood chemistry comparisons between hydrated Tucson and Edmonton mice, Tucson mice had higher chloride (median mmol/L: Tucson: 114.9, Edmonton: 112.5, *p*=0.03, Figure 2a) and creatinine (median mg/dL: Tucson: 0.17, Edmonton: 0.11, *p=*0.02, Figure 2b) levels. Chloride, an electrolyte, is a marker of dehydration and thus expected to be at higher concentrations in the blood of dehydrated animals (MacManes, 2017). Creatinine is a waste product of normal muscle metabolism that is removed from the blood to be excreted by the kidney mainly through glomerular filtration. Blood creatinine levels increase as glomerular filtration, and thus kidney function, decreases and therefore is often used as a measure of kidney health (Kassirer, 1971). The increased levels of serum chloride and creatinine, commonly warnings for declining osmoregulatory function, in healthy Tucson mice suggests that their baseline kidney function differs from that of Edmonton mice and they may be able to function normally despite higher blood osmolarity.

**Figure 2.**
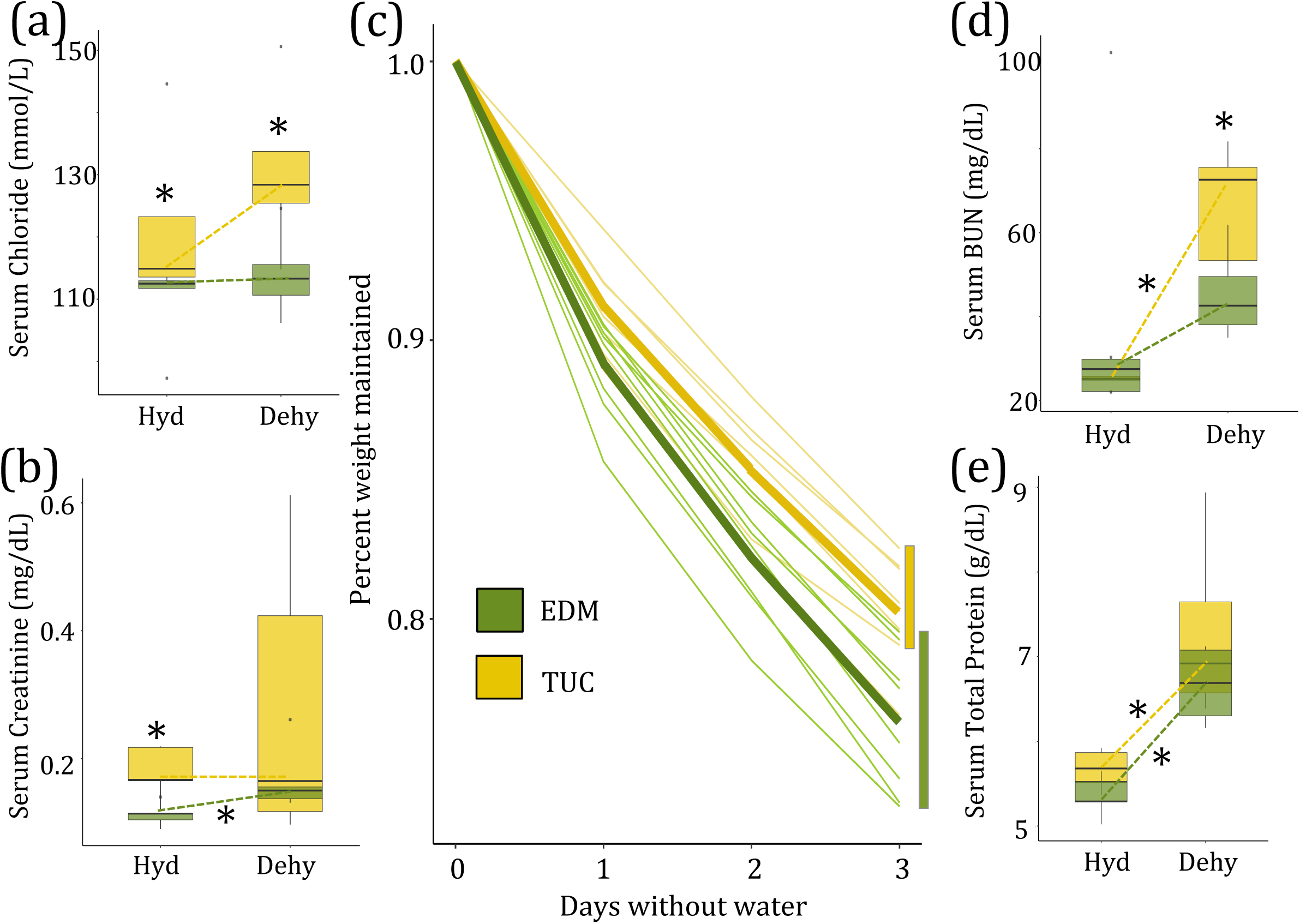
Evolved and plastic phenotypic variation among hydrated and dehydrated mice from desert and non-desert environments. Asterisks denote significant comparisons (p< 0.05) either between lines (Tucson, Edmonton) or between treatments (hydrated, dehydrated). (a) Tucson mice show higher levels of serum chloride than Edmonton mice, both when hydrated and when dehydrated (*p*=0.03 for both). (b) Significant differences between hydrated Edmonton and Tucson mice (*p=*0.02) and between hydrated and dehydrated Edmonton mice (*p*=0.03) in creatinine. (c) Tucson mice show significantly less weight loss after 72 hours of water restriction (*p*=0.027). (d) Significant differences in serum BUN between hydrated and dehydrated Tucson mice (*p* = 0.008) and dehydrated Tucson and Edmonton mice (*p* = 0.03). (e) Significant differences in total protein between hydrated and dehydrated Tucson mice (*p*=0.008) and between hydrated and dehydrated Edmonton mice (*p*=0.01). Vertical lines denote 1.5 * the interquartile range.

Next, we asked how mice from Tucson and mice from Edmonton differed in their response to water stress. We took male full-siblings from the same litter as mice from our hydrated comparison and withheld water from these mice for 72-hours. Hereafter, we refer to the water-restricted group as “dehydrated.” We found that Tucson mice lost significantly less weight than Edmonton mice over the course of 72 hours without access to water (median percent weight maintained: Tucson: 0.82, Edmonton: 0.78, Figure 2c) (Mann-Whitney *U*, *p*=0.027) suggesting that Tucson mice are more buffered against water stress. Comparing blood chemistry measures after mice were subjected to water stress, we found that Tucson mice measured higher in BUN (median mg/dL: Tucson: 64.45, Edmonton: 43.80, *p*=0.03, Figure 2d), chloride (median mmol/L, Tucson: 129.60, Edmonton: 111.95, *p*=0.03, Figure 2a), and potassium (median mmol/L: Tucson: 18.06, Edmonton: 7.78, *p*=0.03, Figure S4a) than Edmonton mice. BUN is a waste product from the liver during the metabolism of protein and increases as glomerular filtration rate and blood volume decreases (Baum et al., 1975). Similarly, high levels of serum potassium often reflect a decrease in filtration of the solute from the blood and thus decreased kidney function (Schwartz, 1955) although this measure is particularly sensitive to lysed blood cells during collection and could reflect the challenge of collecting blood from dehydrated animals. Regardless of treatment, Tucson mice maintained higher serum chloride levels than Edmonton mice. In contrast, potassium levels were higher in Tucson mice only in the dehydrated treatment. High levels of both of these electrolytes are consistent with dehydration. Dehydrated mice from Tucson showed greater indicators for dehydration and kidney dysfunction than Edmonton mice but lost less weight when water stressed. This may reflect a greater evolved capacity to respond to the stress of dehydration. The fact that phenotypic differences in both hydrated and dehydrated animals persist in a common environment indicates that they may have a genetic basis.

While differences between the Tucson and Edmonton lines represent evolved differences, differences between hydrated and dehydrated mice are evidence of plastic responses to water stress. Many of the traits that differed between lines also exhibited phenotypic plasticity in comparisons between hydrated and dehydrated mice within lines. Dehydrated Tucson mice had higher levels of serum BUN compared with hydrated Tucson mice (median mg/dL, Hydrated:25.20, Dehydrated: 64.45, *p*=0.008, Figure 2d). BUN differed both between stressed and control mice from Tucson and was higher than in Edmonton mice when water stressed. Dehydrated Edmonton mice had higher levels of serum creatinine (median mg/dL: Hydrated: 0.11, Dehydrated:0.14, *p*=0.03, Figure 2b) and sodium (mean mmol/L: Hydrated: 153, Dehydrated: 161, *p*=0.01, Figure S4b) compared with hydrated Edmonton mice, reaching the levels for both solutes that were seen in Tucson hydrated and dehydrated animals. The only solute that responded to water stress in both lines was total protein (median g/dL: Edmonton: Hydrated: 5.29, Dehydrated: 6.89, *p*=0.01, Tucson: Hydrated: 5.68, Dehydrated: 7.29, *p*=0.008). Total protein measures the concentration of both albumin and gobulin in the blood which increases with dehydration (Senay & Christensen, 1965). These results indicate that while both lines react physiologically to the stress of dehydration, they may do this through different mechanisms.

### Evolved and plastic transcriptional responses to xeric conditions

Changes in gene expression provide a flexible mechanism for rapidly responding to changes in the local environment, and can also underlie evolutionary divergence. Kidneys are the primary osmoregulatory organ and are essential for homeostasis and solute excretion. To identify candidate genes underlying adaptation to low water environments over short evolutionary timescales as well as plastic responses to water restriction, we sequenced mRNA from kidneys of ten Tucson and ten Edmonton mice, five from the dehydrated and five from the hydrated treatment. Differences in expression between Tucson and Edmonton mice in a common environment represent evolved differences, while differences between dehydrated and hydrated treatments represent a plastic response to water restriction.

We sequenced a total of ∼1.3 billion reads for an average of 26,406,068 uniquely mapped reads per sample which were used to quantify mRNA expression levels and differential expression between samples. Gene expression was measured in a total of 54,233 genes as defined by Ensembl GRCm38 (mm10) with 19,105 genes expressed over a mean of ten reads per sample. Sampling the 1000 genes with the greatest variance, we found that Tucson and Edmonton individuals clustered separately, indicating that more of the gene expression variation was partitioned between line-of-origin than between treatment group (Figure 3a). Edmonton mice clustered into two distinct groups based on treatment (dehydrated vs. hydrated individuals), but dehydrated and hydrated Tucson mice did not form distinct clusters (Figure 3a).

**Figure 3.**
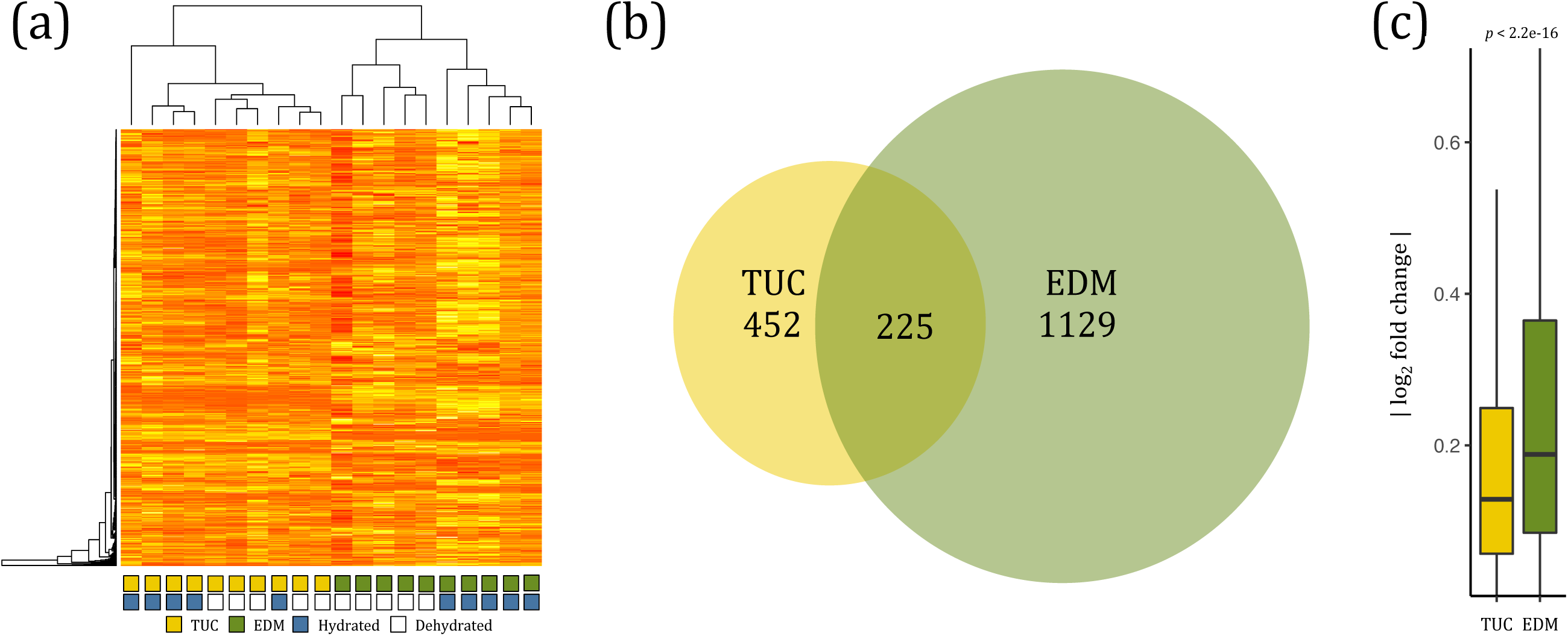
Evolved and plastic gene expression variation among hydrated and dehydrated mice from desert and non-desert environments. (a) Heat map depicting relationships among samples for the top 1000 genes with greatest variance in expression. Expression patterns form two major groups, corresponding to line of origin (Tucson versus Edmonton). Edmonton samples also form clusters based on treatment (hydrated versus dehydrated) while Tucson samples do not. (b) Numbers of differentially expressed genes between dehydrated and hydrated samples in Tucson and Edmonton. Edmonton mice exhibit twice as many genes with differential expression between dehydrated and hydrated conditions compared to Tucson mice. The 225 genes at the intersection represent the shared transcriptional response to water stress. (c) Magnitude of fold changes in each population between dehydrated and hydrated samples. Vertical lines denote 1.5 * the interquartile range.

To identify differential gene expression, we used DESeq2 (Love et al. 2014) to perform pairwise comparisons between: 1) Tucson hydrated *vs.* Edmonton hydrated, and 2) Tucson dehydrated *vs.* Edmonton dehydrated, 3) Tucson dehydrated *vs*. Tucson hydrated, and 4) Edmonton dehydrated *vs.* Edmonton hydrated individuals. Overall, we found more genes were differentially expressed between the two lines than between the treatment groups (FDR of 5%; see methods). A total of 3,935 genes were differentially expressed between hydrated Edmonton and Tucson mice while a total of 3,419 genes were differentially expressed between dehydrated Tucson and Edmonton individuals, a 51% overlap with differences seen between hydrated Tucson and Edmonton mice.

Comparing the hydrated and dehydrated groups within each population (Tucson dehydrated *vs*. Tucson hydrated, and Edmonton dehydrated *vs*. Edmonton hydrated), we found that twice as many genes were differentially expressed in the Edmonton (1354) than in the Tucson comparisons (677 genes) (Chi-square test with Yates Correction, *p* < 0.0001, Figure 3b), with a 225 gene overlap. This 225 gene overlap represents the shared transcriptional response to dehydration with respect to these two lines and is enriched for phenotypes including dehydration (*q*=9×10^-3^) and decreased vasodilation (*q* =3.8×10^-2^) and GO terms involved in regulation of blood pressure (*q* = 4.99×10^-2^). Genes differentially expressed between hydrated and dehydrated Edmonton mice were also enriched for GO terms relevant to water stress, such as renal system processes (*q* = 5.08 × 10^-2^) and regulation of body fluids (*q* = 1.30×10^-2^). Within genes solely differentially expressed between the Tucson groups, we saw enrichment for homeostasis related GO terms (see File S1), but not for any kidney-specific categories. In addition to having a greater number of differentially expressed genes in Edmonton comparisons, we also found that the average magnitude of expression differences between hydrated and dehydrated treatments (|log_2_ fold change|) was significantly greater for Edmonton mice than for Tucson mice (Mann-Whitney *U*, *p* < 2.2 × 10^-16^) (Figure 3c). Together, these results are consistent with Edmonton mice being farther from their physiological optimum when water stressed than are Tucson mice, consistent with the hypothesis that Tucson mice are locally adapted to a water limited environment.

### Dehydrated Edmonton mice show shifts towards Tucson-like expression

Next, we were interested in asking whether the plastic changes in response to water stress in the non-xeric mice (i.e. Edmonton) go in the same or opposite direction as the evolved differences between mice from xeric and non-xeric habitats. Specifically, we were interested in whether Edmonton mice placed under water stress would show transcriptional responses that make them more similar to the base-line condition of Tucson mice. Therefore, we focused on differentially expressed genes between the Edmonton hydrated and dehydrated groups and asked whether the Edmonton dehydrated group showed shifts in expression in the direction of the Tucson hydrated condition (Figure 4a). The majority of these genes (85%, 416 genes) showed changes in the same direction, meaning that the putatively adaptive and plastic responses were in the same direction (+/+ and −/−) (Binomial exact test, *P*<0.0001). Only 15% (74 genes) show changes in opposite directions (+/− and −/+) (Figure 4b).

**Figure 4.**
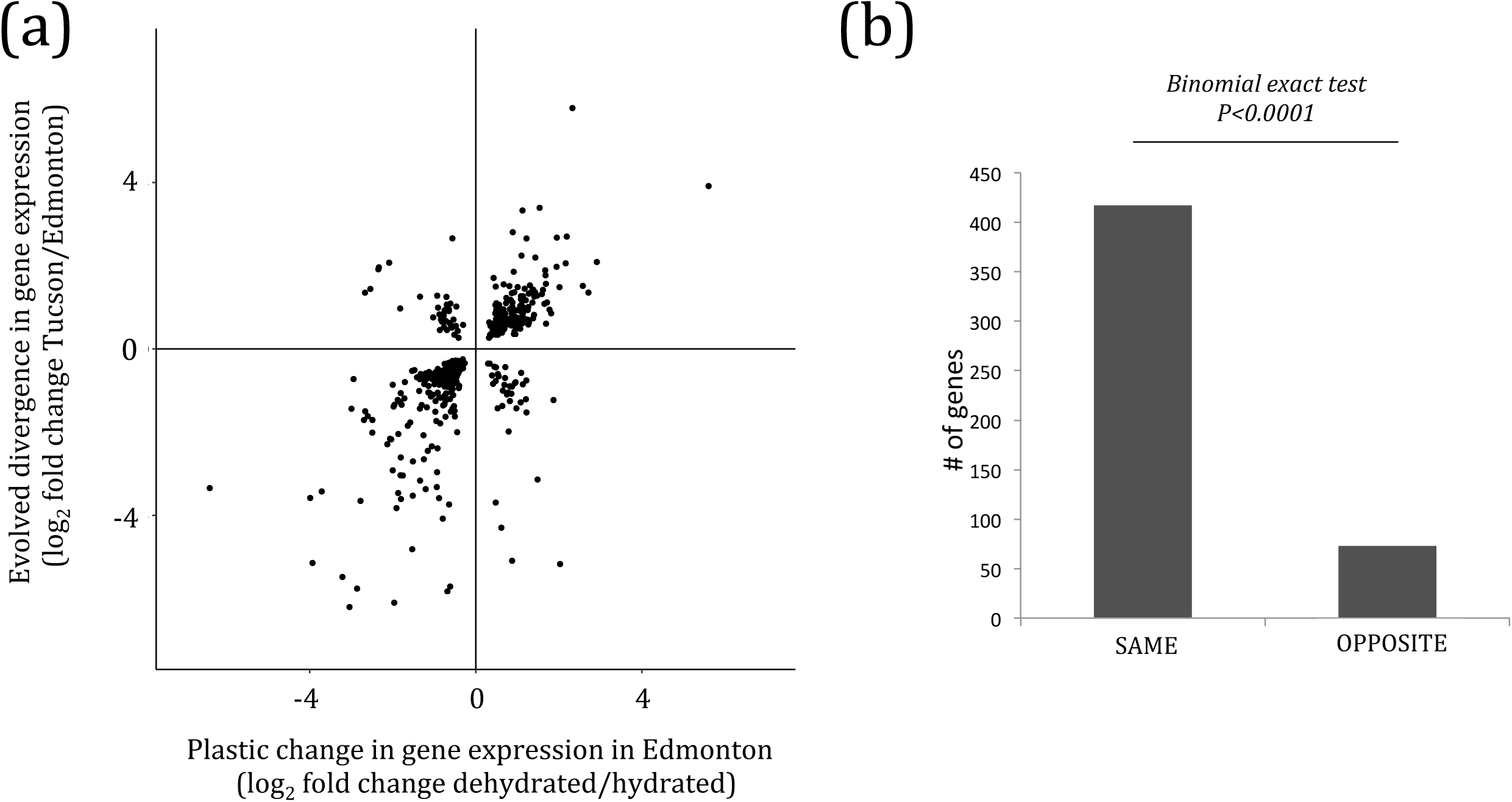
Plastic responses to dehydration in non-desert (Edmonton) mice are mostly in the same direction as evolved differences between non-desert (Edmonton) and desert (Tucson) hydrated mice. (a) Scatterplot comparing the evolved and plastic changes in gene expression. Each point represents a gene and the log2 fold change between Edmonton hydrated vs. dehydrated on the x-axis (plastic response) and Tucson hydrated vs. Edmonton hydrated on the y-axis (evolved divergence). (b) Number of genes in which the evolved and plastic transcriptional responses go in the same direction, and number of genes in which the evolved and plastic transcriptional responses go in the opposite direction.

The group of 416 genes with evolved and plastic changes in the same direction was enriched for GO terms involved in homeostasis and ion transport and for mutant renal/urinary system phenotypes (*q* = 0.035). For example, one of these genes, *Aquaporin 1* (*Aqp1*), showed differences in expression between hydrated and dehydrated Edmonton mice (*q* = 0.0016) and between the hydrated conditions of both lines (*q* = 0.00019)(Figure 5a). In Tucson mice, there was no significant effect of treatment on expression level, but in Edmonton mice, expression increased in response to water stress recapitulating the constitutive expression level of the mice from Tucson. Aquaporins are a family of membrane proteins which form channels used to transport water and small solutes across cell membranes. *Aqp1* is expressed in the descending loop of Henle, and channels formed from this protein are the main route through which water is reabsorbed in this region (Chou et al., 1999). It is known to affect urine concentrating ability, response to dehydration, and renal water transport in lab lines of house mice (Ma et al., 1998; Sohara et al., 2005) and has been identified in a several studies related to desert adaptation in rodents (reviewed in Pannabecker, 2015). In our analyses, expression of *Aqp1* was associated with variation in six of the nine measured phenotypes (Creatinine, *p*=0.020; total protein, *p*=0.012; potassium, *p*=0.0094; weight loss, *p*=0.016; kidney weight, *p*=0.0077; RWC, p=0.022).

**Figure 5.**
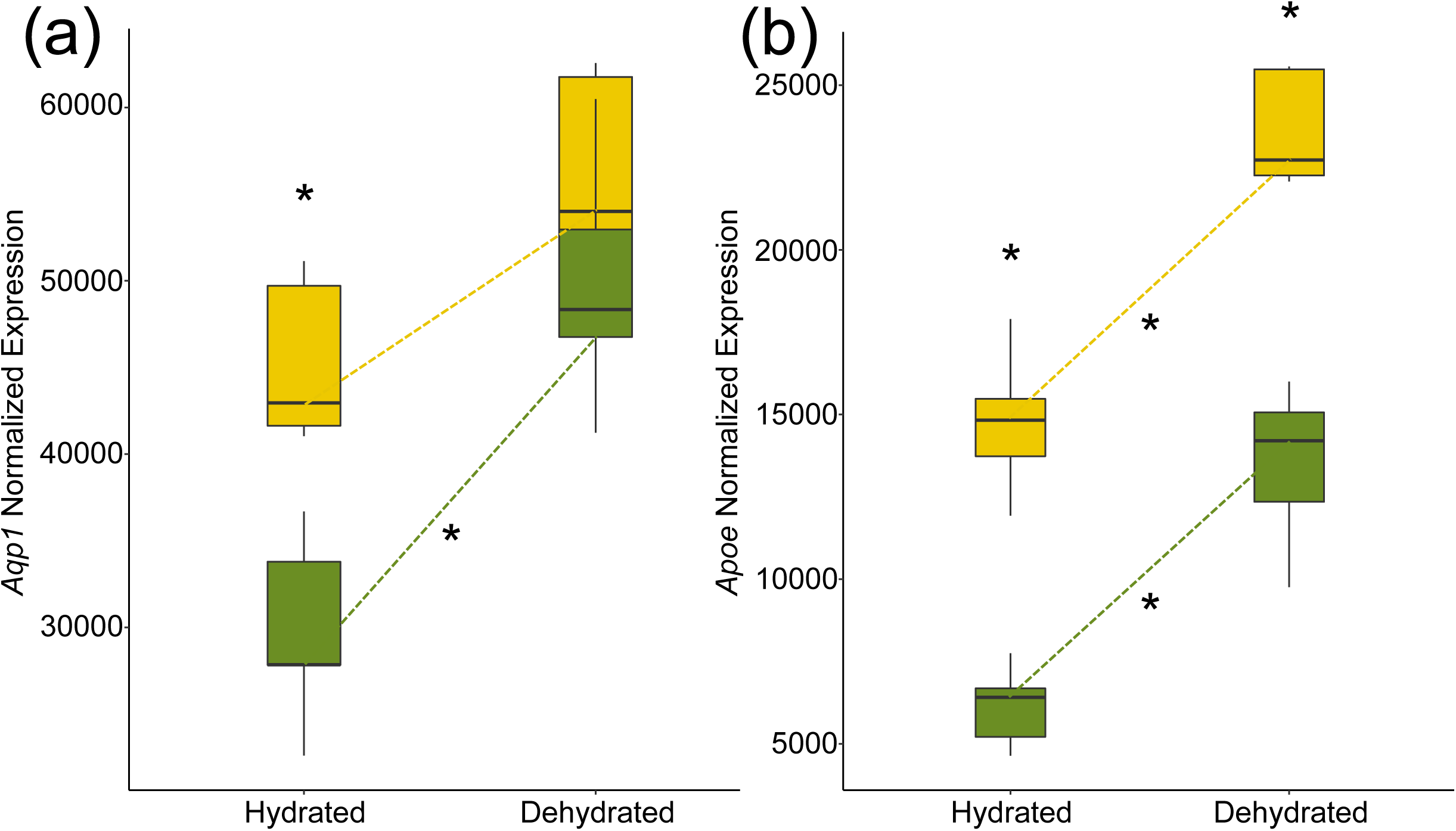
Expression variation at two genes that may underlie adaptation to a desert environment. Normalized gene expression in hydrated and dehydrated mice from Tucson (gold) and Edmonton (green). Asterisks denote significant comparisons (p< 0.05) either between lines (Tucson, Edmonton) or between treatments (hydrated, dehydrated). (a) *Aqp1*. (b) *Apoe*. For both genes, the dehydrated Edmonton mice recapitulate the baseline expression level seen in hydrated Tucson mice. For *Aqp1*, the plastic response is attenuated in Tucson mice (a), while for *Apoe*, it is not (b). Vertical lines denote 1.5 * the interquartile range.

Of the 416 genes where dehydrated Edmonton mice showed shifts towards Tucson-like expression, the majority (87%) were not differentially expressed between hydrated and dehydrated Tucson mice. For the 54 of these genes that were differentially expressed in the Tucson comparison, in all cases the plastic response was observed to be changing in the same direction as in Edmonton mice.

### Co-expressed sets of genes are associated with phenotypic variation in Tucson and Edmonton mice

To identify transcriptional networks associated with phenotypic variation in Tucson and Edmonton mice, we performed a weighted gene co-expression network analysis (WGCNA). This analysis uncovers genes with highly correlated expression profiles and groups them into modules reflecting hypotheses about connectivity. We identified 54 co-expression modules, with at least one module significantly associated with each of the nine phenotypes (Figure S5). Of these, the “salmon” module (Figure S6a) was of particular interest because it was significantly associated with all nine phenotypes (BUN, *p*=0.02; Creatinine, *p*=0.006; total protein, *p* = 1 × 10^-4^; Chloride, *p* = 0.002; Potassium, *p* = 0.001; Sodium level, *p* = 0.01; weight loss, *p* = 0.007; kidney weight, *p* = 0.03; RWC, p= 0.006) as well as population of origin (*p* = 0.006) and treatment (*p* = 0.001). Genes in this module were enriched for several metabolic processes, including glyceraldehyde-3-phosphate metabolic process (*q* = 6.85 × 10^-9^), ribose phosphate metabolic process (*q* = 1.45 × 10^-8^), and carbohydrate derivative metabolic process (*q* = 1.3 × 10^-7^). Visualizing the most connected genes in the salmon module, we identified *Tmtc1*, *Apoe*, and *Sult1a1* as the most centrally located hub genes (Figure S6a). All three of these genes were identified in our previous analysis as genes for which dehydrated Edmonton mice showed shifts towards Tucson-like expression (Figure 4). *Apoe* is of particular interest because of its documented role in kidney function. It is thought to play an important role in renal damage protection (Bonomini et al., 2011; Wen et al., 2002), and laboratory mutants show a number physiological and morphological changes similar to kidney disease phenotypes (Bonomini et al., 2011). A*poe* expression responded in the same direction and in a similar magnitude in both lines under water stress. Interestingly, under water stress, the expression level in Edmonton recapitulated the constitutive expression level of hydrated Tucson mice (Figure 5b). *Apoe* expression was also correlated with eight of the nine measured phenotypes (Creatinine, *p*=0.034; total protein, *p*=0.00077; chloride, *p*=0.0055; potassium, p=0.0041; sodium, *p*=0.034; weight loss, *p*=0.014; kidney weight, *p*=0.025; RWC, *p*=0.0036).

In addition to shifts in the expression of entire co-expression modules, populations may differ as a consequence of altered co-expression between pairs of genes, called differential co-expression. In order to identify genes that show differential co-expression between Tucson and Edmonton, we calculated the average change in correlation between the two lines across all gene pairs using the program DGCA (Differential Gene Correlation Analysis) (McKenzie et al., 2016). We identified 182 genes that showed significantly different co-expression between Tucson and Edmonton individuals (*q* <0.10, see methods). These genes were enriched for several GO categories including glycolysis: generation of precursor metabolites and energy (*q* = 0.0013), cellular amino acid metabolic process (*q* = 0.00046), and lipid metabolic process (*q* = 0.0014). This gene set was also enriched for mutant phenotypes related to renal/urinary system (*q* = 0.00001), abnormal urine homeostasis (*q* = 0.005), and abnormal ion homeostasis (*q* = 0.01). One of the genes with significant changes in co-expression was *Aqp1*, which is involved in renal water transport. We also found that several solute carriers (*Slc22a19, Slc11a2, Slc36a1, Slc47a1, Slc12a1, Slco3a1*) showed evidence of differential co-expression, including one (*Slc47a1*) that has been shown to be under positive selection in the desert-adapted cactus mouse, *Peromyscus eremicus* (Kordonowy et al., 2017). *Slc47a1* mouse mutants are also associated with increased BUN, increased circulating creatinine level, and kidney degeneration (Tsuda et al., 2009). Altogether, these results suggest that Tucson and Edmonton mice show shifts in the expression of co-expression modules as well as changes in the co-expression associations between sets of genes.

## Discussion

Here we have described phenotypic and transcriptional divergence between descendants of mice from a desert environment and descendants of mice from a more mesic environment when reared under identical laboratory conditions. We also described plastic responses in these mice under conditions of water stress. First, we showed that inbred lines derived from the Sonoran desert consume less water than do mice from other populations in the Americas. Next, comparisons between a single line from Tucson and a single line from Edmonton revealed many phenotypic differences in a common environment, both at the organismal level and at the gene expression level in the kidney. The fact that these differences were present after multiple generations in the lab indicates that they are genetically based. Nonetheless, these same traits reveal considerable plasticity in comparisons between control mice and mice under conditions of water stress. Notably, we found that Tucson mice showed attenuated responses to water stress. After a 72 hour period without water, Tucson mice lost less weight and showed fewer expression differences in the kidney. Surprisingly, the blood chemistry of Tucson mice was consistent with higher levels of dehydration and reduced kidney function, both under standard and water-restricted conditions. However, phenotypes that appear maladaptive in one genomic or environmental context may be adaptive in another (e.g., Riddle et al., 2018). Altogether, these results are consistent with genetic changes to Tucson mice following their invasion of the desert environment.

Phenotypic plasticity may allow animals to persist in harsh new environments if the plastic responses bring individuals closer to the local optimum (reviewed by Ghalambor et al., 2007). Adaptive plasticity can be followed by genetic changes as populations become established, in a process called “genetic assimilation” or the “Baldwin Effect” (Price et al., 2003; Robinson & Dukas, 1999; Simpson, 1953; Waddington, 1942, 1952, 1953). We found that the evolved differences in kidney gene expression between Tucson and Edmonton mice were generally in the same direction as plastic changes in Edmonton individuals under water stress. Consequently, under water stress, gene expression in Edmonton mice becomes more similar to that of Tucson mice under hydrated conditions. This result is consistent with the idea that plastic responses to short-term water stress are an example of adaptive plasticity. Thus, plastic responses to water stress may have helped facilitate the colonization and subsequent adaptation of house mice to the desert environment.

Our finding that plastic responses generally go in the same direction as evolved responses stands in contrast to several recent studies. For example, an allele of the *Epas1* gene confers adaptation to high altitude in Tibetans by attenuating the maladaptive plastic response of increased hemoglobin concentration (Beall et al., 2010; Jeong et al., 2018; Simonson et al., 2010; Yi et al., 2010). Similarly, most gene expression changes in the brains of guppies reared in the absence of predators go in the opposite direction of those seen in populations that have evolved without predators (Ghalambor et al. 2015). In both cases, the selective agent (hypoxia in humans or absence of predators in fish) may have not been present in the recent history of the populations exhibiting non-adaptive plasticity. In contrast, we speculate that occasional periods of water stress are probably common in many populations of mice, including in places, like Edmonton, that are not in deserts. Under such situations, selection is expected to favor an adaptive plastic response.

While adaptive plasticity may facilitate the colonization of new environments, it can also slow or impede adaptive evolution if, by moving individuals closer to the optimum, genetic variation is shielded from natural selection (Ghalambor et al., 2007; Price et al., 2003). However, when plastic responses to a new environment are incomplete, directional selection may favor a more extreme phenotype and lead to subsequent genetic changes (Price et al. 2003). While all house mice, even those in mesic environments, likely undergo short periods of water stress, water stress is likely to be more severe in desert environments. Phenotypic comparisons between Edmonton and Tucson mice suggest that plastic responses to water stress in Edmonton mice may be suboptimal. Tucson mice drink less water on average and lose less weight in response to dehydration, indicating that these animals are buffered against water stress in a way that Edmonton mice are not. The blood chemistry comparisons reported here also suggest there are differences between Tucson and Edmonton kidney function and homeostasis. Consequently, we suggest that while phenotypic plasticity likely helped house mice colonize the Sonoran desert, the xeric environment still imposed sufficient selective pressure for subsequent genetic changes.

Finally, the comparison of gene expression changes both between treatments and between lines has identified a few genes that may be important in the adaptive response. Expression changes pinpoint a number of interesting candidates, including *Aqp1* and *Apoe*. Future studies aimed at identifying *cis-*regulatory variation at these genes might help to pinpoint causative mutations underlying adaptation to desert environments.

## Acknowledgements

We thank members of the Nachman Lab for valuable comments and discussions. We thank Taichi Suzuki, Megan Phifer-Rixey, Felipe Martins, Dana Lin, Michael Sheehan, and Mallory Ballinger for help with animal husbandry or for collecting the wild mice used to establish the lines for this study. We thank Lydia Smith of UC Berkeley and Eugene Dunn of UC Davis for their technical expertise. This work was facilitated by an Extreme Science and Engineering Discovery Environment (XSEDE) allocation to M.W.N. XSEDE is supported by National Science Foundation (NSF) grant number ACI-1548562. This work was supported by an NIH grant to MWN (R01 GM127468) and a NSF Doctoral Dissertation Improvement Grant to NKJB (1601827).

## Data Accessibility Statement

Illumina sequencing data from this study will be submitted to the NCBI BioProject (https://www.ncbi.nlm.nih.gov/bioproject) under accession number xxxxxxx.

## Author Contributions

This study was designed by all authors. NKJB conducted the experiments, and NKJB and KLM analyzed the data. The paper was written by NKJB and edited by MWN and KLM.

## References

Al-kahtani, M. A., Zuleta, C., Caviedes-Vidal, E., & Garland, Jr., Theodore. (2004). Kidney Mass and Relative Medullary Thickness of Rodents in Relation to Habitat, Body Size, and Phylogeny. Physiological and Biochemical Zoology: Ecological and Evolutionary Approaches, 77(3), 346–365. https://doi.org/10.1086/420941

Anders, S., Pyl, P. T., & Huber, W. (2015). HTSeq—A Python framework to work with high-throughput sequencing data. Bioinformatics, 31(2), 166–169. https://doi.org/10.1093/bioinformatics/btu638

Baldwin, J. M. (1896). A New Factor in Evolution. 11.

Baum, N., Dichoso, C. C., & Carlton, C. E. (1975). Blood urea nitrogen and serum creatinine: Physiology and interpretations. Urology, 5(5), 583–588. https://doi.org/10.1016/0090-4295(75)90105-3

Beall, C. M., Cavalleri, G. L., Deng, L., Elston, R. C., Gao, Y., Knight, J., Li, C., Li, J. C., Liang, Y., McCormack, M., Montgomery, H. E., Pan, H., Robbins, P. A., Shianna, K. V., Tam, S. C., Tsering, N., Veeramah, K. R., Wang, W., Wangdui, P., … Zheng, Y. T. (2010). Natural selection on EPAS1 (HIF2α) associated with low hemoglobin concentration in Tibetan highlanders. Proceedings of the National Academy of Sciences, 107(25), 11459–11464. https://doi.org/10.1073/pnas.1002443107

Beuchat, C. A. (1990). Body size, medullary thickness, and urine concentrating ability in mammals. American Journal of Physiology-Regulatory, Integrative and Comparative Physiology, 258(2), R298–R308. https://doi.org/10.1152/ajpregu.1990.258.2.R298

Bolger, A. M., Lohse, M., & Usadel, B. (2014). Trimmomatic: A flexible trimmer for Illumina sequence data. Bioinformatics, 30(15), 2114–2120. https://doi.org/10.1093/bioinformatics/btu170

Bonomini, F., Rodella, L. F., Moghadasian, M., Lonati, C., Coleman, R., & Rezzani, R. (2011). Role of apolipoprotein E in renal damage protection. Histochemistry and Cell Biology, 135(6), 571–579. https://doi.org/10.1007/s00418-011-0815-1

Chou, C.-L., Knepper, M. A., Hoek, A. N. van, Brown, D., Yang, B., Ma, T., & Verkman, A. S. (1999). Reduced water permeability and altered ultrastructure in thin descending limb of Henle in aquaporin-1 null mice. The Journal of Clinical Investigation, 103(4), 491–496. https://doi.org/10.1172/JCI5704

Corl, A., Bi, K., Luke, C., Challa, A. S., Stern, A. J., Sinervo, B., & Nielsen, R. (2018). The Genetic Basis of Adaptation following Plastic Changes in Coloration in a Novel Environment. Current Biology, 28(18), 2970–2977.e7. https://doi.org/10.1016/j.cub.2018.06.075

Dobin, A., Davis, C. A., Schlesinger, F., Drenkow, J., Zaleski, C., Jha, S., Batut, P., Chaisson, M., & Gingeras, T. R. (2013). STAR: Ultrafast universal RNA-seq aligner. Bioinformatics, 29(1), 15–21. https://doi.org/10.1093/bioinformatics/bts635

Donald, J., & Pannabecker, T. L. (2015). Osmoregulation in Desert-Adapted Mammals. In K. A. Hyndman & T. L. Pannabecker (Eds.), Sodium and Water Homeostasis: Comparative, Evolutionary and Genetic Models (pp. 191–211). Springer. https://doi.org/10.1007/978-1-4939-3213-9_10

Eden, E., Navon, R., Steinfeld, I., Lipson, D., & Yakhini, Z. (2009). GOrilla: A tool for discovery and visualization of enriched GO terms in ranked gene lists. BMC Bioinformatics, 10(1), 48. https://doi.org/10.1186/1471-2105-10-48

Ghalambor, C. K., McKAY, J. K., Carroll, S. P., & Reznick, D. N. (2007). Adaptive versus non-adaptive phenotypic plasticity and the potential for contemporary adaptation in new environments. Functional Ecology, 21(3), 394–407. https://doi.org/10.1111/j.1365-2435.2007.01283.x

Ghalambor, Cameron K., Hoke, K. L., Ruell, E. W., Fischer, E. K., Reznick, D. N., & Hughes, K. A. (2015). Non-adaptive plasticity potentiates rapid adaptive evolution of gene expression in nature. Nature, 525(7569), 372–375. https://doi.org/10.1038/nature15256

Giorello, F. M., Feijoo, M., D’Elía, G., Naya, D. E., Valdez, L., Opazo, J. C., & Lessa, E. P. (2018). An association between differential expression and genetic divergence in the Patagonian olive mouse (Abrothrix olivacea). Molecular Ecology, 27(16), 3274–3286. https://doi.org/10.1111/mec.14778

Haines, H., Ciskowski, C., & Harms, V. (1973). Acclimation to Chronic Water Restriction in the Wild House Mouse Mus musculus. Physiological Zoology, 46(2), 110–128. https://doi.org/10.1086/physzool.46.2.30155592

Haines, H., & Schmidt-Nielsen, K. (1967). Water Deprivation in Wild House Mice. Physiological Zoology, 40(4), 424–431. https://doi.org/10.1086/physzool.40.4.30158460

Huang, Y., & Agrawal, A. F. (2016). Experimental Evolution of Gene Expression and Plasticity in Alternative Selective Regimes. PLOS Genetics, 12(9), e1006336. https://doi.org/10.1371/journal.pgen.1006336

Jeong, C., Witonsky, D. B., Basnyat, B., Neupane, M., Beall, C. M., Childs, G., Craig, S. R., Novembre, J., & Rienzo, A. D. (2018). Detecting past and ongoing natural selection among ethnically Tibetan women at high altitude in Nepal. PLOS Genetics, 14(9), e1007650. https://doi.org/10.1371/journal.pgen.1007650

Kassirer, J. P. (1971). Clinical Evaluation of Kidney Function: Glomerular Function. New England Journal of Medicine, 285(7), 385–389. https://doi.org/10.1056/NEJM197108122850706

Koford, C. B. (1968). Peruvian Desert Mice: Water Independence, Competition, and Breeding Cycle near the Equator. Science, 160(3827), 552–553. https://doi.org/10.1126/science.160.3827.552

Kordonowy, L., Lombardo, K. D., Green, H. L., Dawson, M. D., Bolton, E. A., LaCourse, S., & MacManes, M. D. (2017). Physiological and biochemical changes associated with acute experimental dehydration in the desert adapted mouse, Peromyscus eremicus. Physiological Reports, 5(6). https://doi.org/10.14814/phy2.13218

Langfelder, P., & Horvath, S. (2008). WGCNA: An R package for weighted correlation network analysis. BMC Bioinformatics, 9(1), 559. https://doi.org/10.1186/1471-2105-9-559

Levis, N. A., & Pfennig, D. W. (2016). Evaluating ‘Plasticity-First’ Evolution in Nature: Key Criteria and Empirical Approaches. Trends in Ecology & Evolution, 31(7), 563–574. https://doi.org/10.1016/j.tree.2016.03.012

Love, M. I., Huber, W., & Anders, S. (2014). Moderated estimation of fold change and dispersion for RNA-seq data with DESeq2. Genome Biology, 15(12), 550. https://doi.org/10.1186/s13059-014-0550-8

Lynch, C. B. (1992). Clinal Variation in Cold Adaptation in Mus domesticus: Verification of Predictions from Laboratory Populations. The American Naturalist, 139(6), 1219–1236. https://doi.org/10.1086/285383

Ma, T., Yang, B., Gillespie, A., Carlson, E. J., Epstein, C. J., & Verkman, A. S. (1998). Severely Impaired Urinary Concentrating Ability in Transgenic Mice Lacking Aquaporin-1 Water Channels. Journal of Biological Chemistry, 273(8), 4296–4299. https://doi.org/10.1074/jbc.273.8.4296

Mack, K. L., Ballinger, M. A., Phifer-Rixey, M., & Nachman, M. W. (2018). Gene regulation underlies environmental adaptation in house mice. Genome Research, 28(11), 1636–1645. https://doi.org/10.1101/gr.238998.118

MacManes, M. D. (2017). Severe acute dehydration in a desert rodent elicits a transcriptional response that effectively prevents kidney injury. American Journal of Physiology-Renal Physiology, 313(2), F262–F272. https://doi.org/10.1152/ajprenal.00067.2017

Marra, N. J., Romero, A., & DeWoody, J. A. (2014). Natural selection and the genetic basis of osmoregulation in heteromyid rodents as revealed by RNA-seq. Molecular Ecology, 23(11), 2699–2711. https://doi.org/10.1111/mec.12764

McKenzie, A. T., Katsyv, I., Song, W.-M., Wang, M., & Zhang, B. (2016). DGCA: A comprehensive R package for Differential Gene Correlation Analysis. BMC Systems Biology, 10(1), 106. https://doi.org/10.1186/s12918-016-0349-1

Pannabecker, T. L. (2015). Aquaporins in Desert Rodent Physiology. The Biological Bulletin, 229(1), 120–128. https://doi.org/10.1086/BBLv229n1p120

Phifer-Rixey, M., Bi, K., Ferris, K. G., Sheehan, M. J., Lin, D., Mack, K. L., Keeble, S. M., Suzuki, T. A., Good, J. M., & Nachman, M. W. (2018). The genomic basis of environmental adaptation in house mice. PLOS Genetics, 14(9), e1007672. https://doi.org/10.1371/journal.pgen.1007672

Phifer-Rixey, M., & Nachman, M. W. (2015). Insights into mammalian biology from the wild house mouse Mus musculus. ELife, 4, e05959. https://doi.org/10.7554/eLife.05959

Price, T. D., Qvarnström, A., & Irwin, D. E. (2003). The role of phenotypic plasticity in driving genetic evolution. Proceedings of the Royal Society of London. Series B: Biological Sciences, 270(1523), 1433–1440. https://doi.org/10.1098/rspb.2003.2372

Riddle, M. R., Aspiras, A. C., Gaudenz, K., Peuß, R., Sung, J. Y., Martineau, B., Peavey, M., Box, A. C., Tabin, J. A., McGaugh, S., Borowsky, R., Tabin, C. J., & Rohner, N. (2018). Insulin resistance in cavefish as an adaptation to a nutrient-limited environment. Nature, 555(7698), 647–651. https://doi.org/10.1038/nature26136

Robinson, B. W., & Dukas, R. (1999). The Influence of Phenotypic Modifications on Evolution: The Baldwin Effect and Modern Perspectives. Oikos, 85(3), 582–589. JSTOR. https://doi.org/10.2307/3546709

Schwartz, W. B. (1955). Potassium and the Kidney. New England Journal of Medicine, 253(14), 601–608. https://doi.org/10.1056/NEJM195510062531405

Senay, L. C., & Christensen, M. L. (1965). Changes in blood plasma during progressive dehydration. Journal of Applied Physiology, 20(6), 1136–1140. https://doi.org/10.1152/jappl.1965.20.6.1136

Simonson, T. S., Yang, Y., Huff, C. D., Yun, H., Qin, G., Witherspoon, D. J., Bai, Z., Lorenzo, F. R., Xing, J., Jorde, L. B., Prchal, J. T., & Ge, R. (2010). Genetic Evidence for High-Altitude Adaptation in Tibet. Science, 329(5987), 72–75. https://doi.org/10.1126/science.1189406

Simpson, G. G. (1953). The Baldwin Effect. Evolution, 7(2), 110–117. https://doi.org/10.1111/j.1558-5646.1953.tb00069.x

Sohara, E., Rai, T., Miyazaki, J., Verkman, A. S., Sasaki, S., & Uchida, S. (2005). Defective water and glycerol transport in the proximal tubules of AQP7 knockout mice. American Journal of Physiology-Renal Physiology, 289(6), F1195–F1200. https://doi.org/10.1152/ajprenal.00133.2005

Tsuda, H., Isaka, Y., Takahara, S., & Horio, M. (2009). Discrepancy between serum levels of low molecular weight proteins in acute kidney injury model rats with bilateral ureteral obstruction and bilateral nephrectomy. Clinical and Experimental Nephrology, 13(6), 567–570. https://doi.org/10.1007/s10157-009-0203-5

Waddington, C. H. (1942). Canalization of Development and the Inheritance of Acquired Characters. Nature, 150(3811), 563–565. https://doi.org/10.1038/150563a0

Waddington, C. H. (1952). Selection of the Genetic Basis for an Acquired Character. Nature, 170(4315), 71–71. https://doi.org/10.1038/170071a0

Waddington, C. H. (1953). Genetic Assimilation of an Acquired Character. Evolution, 7(2), 118–126. https://doi.org/10.1111/j.1558-5646.1953.tb00070.x

Wen, M., Segerer, S., Dantas, M., Brown, P. A., Hudkins, K. L., Goodpaster, T., Kirk, E., LeBoeuf, R. C., & Alpers, C. E. (2002). Renal Injury in Apolipoprotein E–Deficient Mice. Laboratory Investigation, 82(8), 999–1006. https://doi.org/10.1097/01.LAB.0000022222.03120.D4

Weng, M.-P., & Liao, B.-Y. (2017). modPhEA: Model organism Phenotype Enrichment Analysis of eukaryotic gene sets. Bioinformatics, 33(21), 3505–3507. https://doi.org/10.1093/bioinformatics/btx426

Wu, H., Guang, X., Al-Fageeh, M. B., Cao, J., Pan, S., Zhou, H., Zhang, L., Abutarboush, M. H., Xing, Y., Xie, Z., Alshanqeeti, A. S., Zhang, Y., Yao, Q., Al-Shomrani, B. M., Zhang, D., Li, J., Manee, M. M., Yang, Z., Yang, L., … Wang, J. (2014). Camelid genomes reveal evolution and adaptation to desert environments. Nature Communications, 5(1), 1–10. https://doi.org/10.1038/ncomms6188

Yi, X., Liang, Y., Huerta-Sanchez, E., Jin, X., Cuo, Z. X. P., Pool, J. E., Xu, X., Jiang, H., Vinckenbosch, N., Korneliussen, T. S., Zheng, H., Liu, T., He, W., Li, K., Luo, R., Nie, X., Wu, H., Zhao, M., Cao, H., … Wang, J. (2010). Sequencing of 50 Human Exomes Reveals Adaptation to High Altitude. Science, 329(5987), 75–78. https://doi.org/10.1126/science.1190371

