## Supplemental Tables 1-6 for "Plasticity in gene expression facilitates invasion of the desert environment in house mice"

MOLECULAR ECOLOGY

### Table of Contents:


| | |
|---|---|
| **Figure S1** | Page 2 |
| **Figure S2** | Page 3 |
| **Figure S3** | Page 4 |
| **Figure S4** | Page 5 |
| **Figure S5** | Page 6 |
| **Figure S6** | Page 7 |

MOLECULAR ECOLOGY

**Figure S1**. Relative water consumption in hydrated control mice from different environments. Edmonton mice consume significantly more water adjusted for body mass than do Tucson mice (*$p$ = 0.0026). Vertical lines denote 1.5 * the interquartile range.

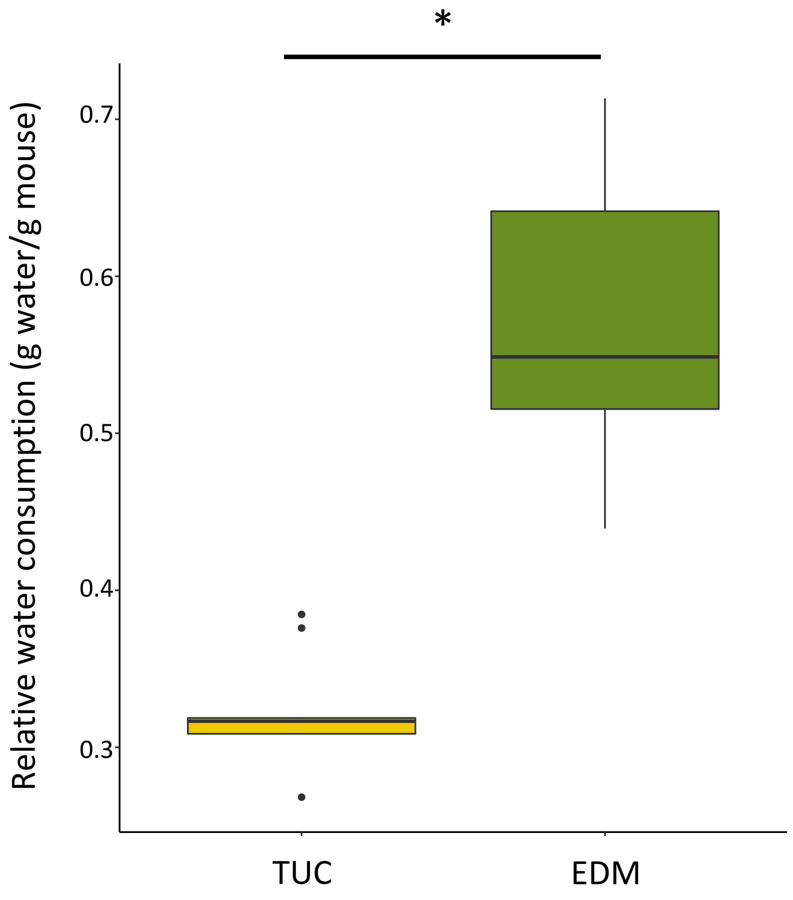

MOLECULAR ECOLOGY

**Figure S2.** Relative kidney weight in hydrated control mice from different environments. Edmonton mice have heavier kidneys relative to their body weight than Tucson mice (*$p$ = 0.00015). Vertical lines denote 1.5 * the interquartile range.

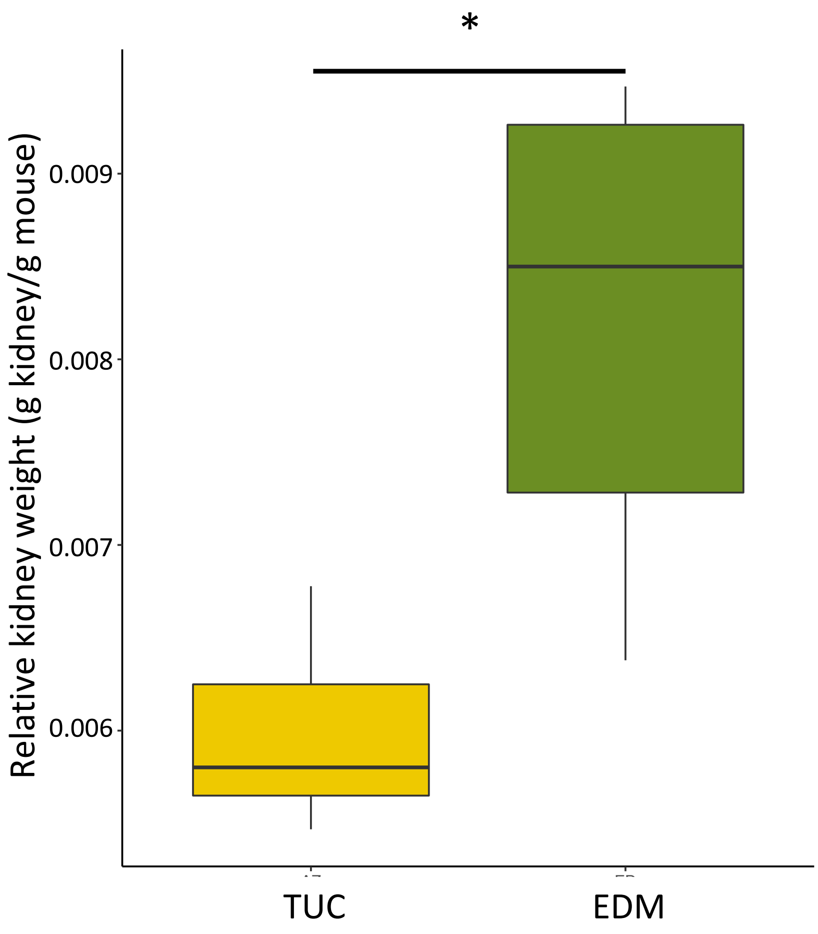

MOLECULAR ECOLOGY

**Figure S3.** Kidney histology from hydrated control mice from different environments. Tucson and Edmonton mice do not show significant differences in (a) renal cortex thickness ($p$ = 0.15) nor (b) the ratio of renal papilla thickness to cortical thickness ($p$ = 0.31). Vertical lines denote 1.5 * the interquartile range. (c) Example cross-section with measurements of a Tucson kidney. (d) Example cross-section with measurements of an Edmonton kidney.

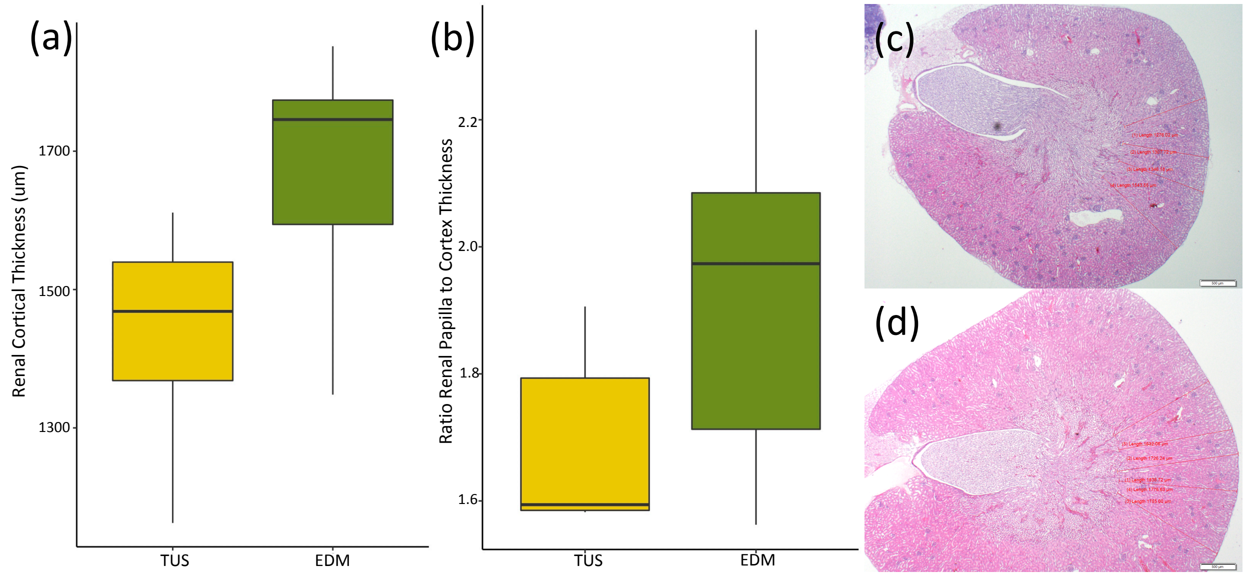

MOLECULAR ECOLOGY

**Figure S4**. Evolved and plastic phenotypic variation among hydrated and dehydrated mice from desert and non-desert environments. Asterisks denote significant comparisons ($p < 0.05$) either between lines (Tucson, Edmonton) or between treatments (hydrated, dehydrated). (a) Serum potassium concentration is significantly different between dehydrated Tucson mice and dehydrated Edmonton mice (median mmol/L: Tucson: 7.78, Edmonton: 18.06, $p$=0.32). (b) Serum sodium concentration is significantly different between hydrated and dehydrated Edmonton mice (median mmol/L: Hydrated: 153, Dehydrated: 161, $p$=0.01). Vertical lines denote 1.5 * the interquartile range.

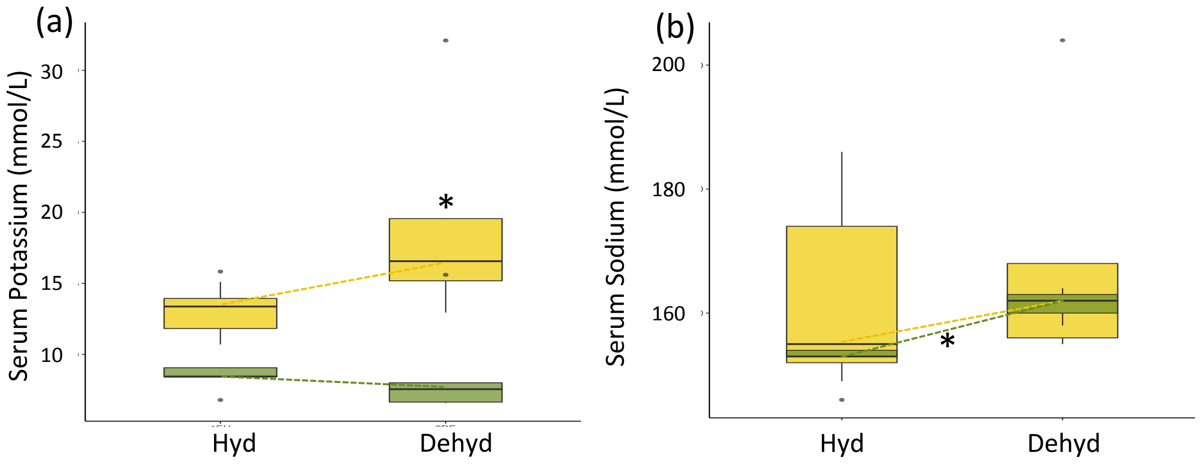

MOLECULAR ECOLOGY

**Figure S5**. Results from a weighted gene co-expression analysis (WGCNA). Modules (rows represented by colors in the first column) represent groups of genes with highly correlated expression values. Each module is represented by the significance of its association with each phenotype, population, or treatment in each column. The "salmon" module is significantly associated with all nine parameters of interest.

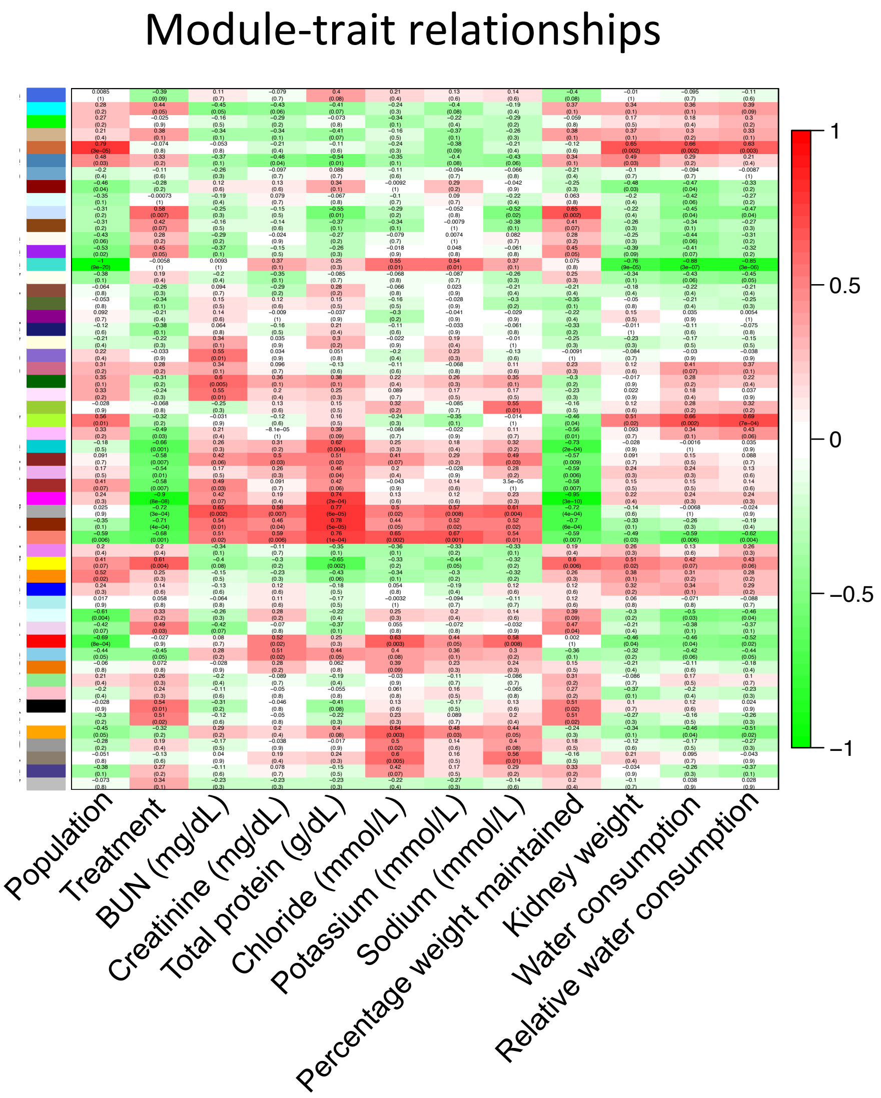

MOLECULAR ECOLOGY

**Figure S6.** Network of genes identified by WGCNA in the "salmon" module. Genes with the greatest number of connections are of particular interest including *Apoe* which is associated with kidney-related phenotypes.

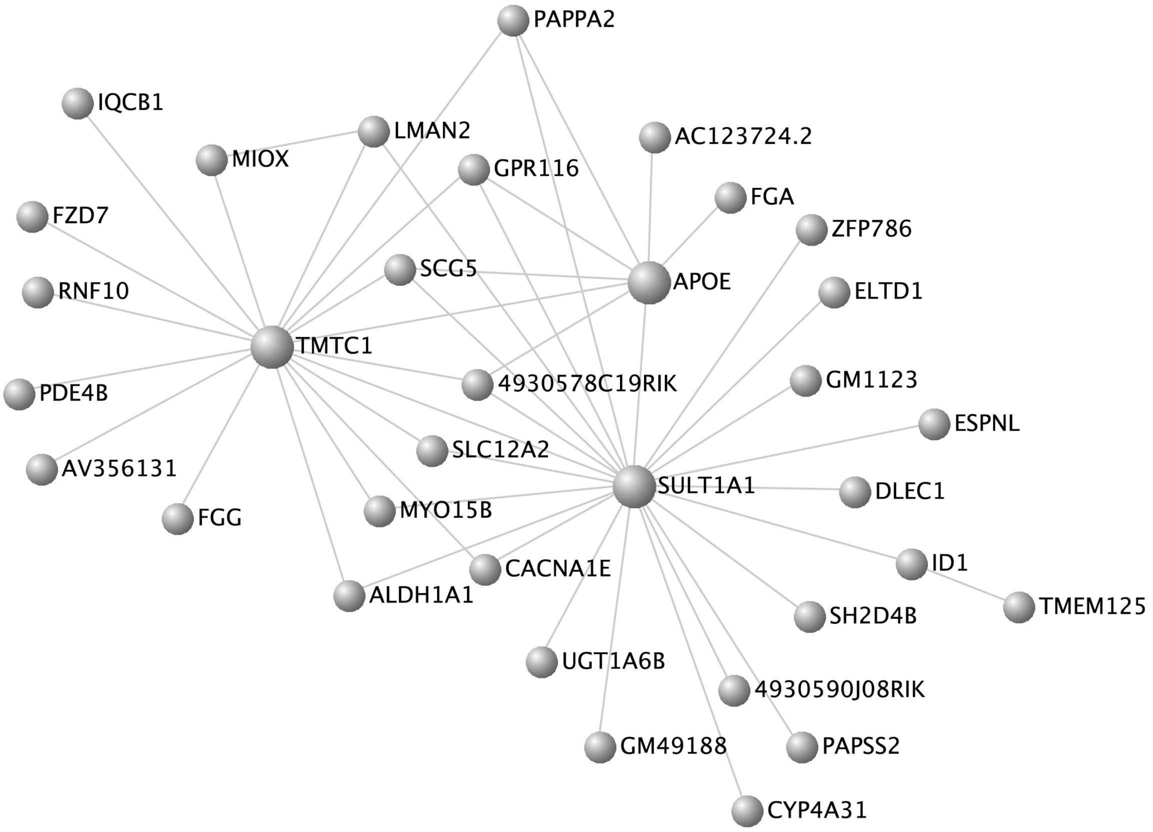
